# Competence of *Corynebacterium glutamicum* as a host for the production of type I polyketides

**DOI:** 10.1101/622399

**Authors:** Nicolai Kallscheuer, Hirokazu Kage, Lars Milke, Markus Nett, Jan Marienhagen

**Affiliations:** Institute of Bio- and Geosciences, IBG-1: Biotechnology, Forschungszentrum Jülich GmbH, D-52425 Jülich, Germany; TU Dortmund University, Department of Biochemical and Chemical Engineering, D-44227 Dortmund, Germany; Institute of Biotechnology, RWTH Aachen University, Worringer Weg 3, D-52074 Aachen, Germany

**Author notes:** Corresponding author Dr. Jan Marienhagen.

**Keywords:** polyketide, 6-methylsalicylate, type I polyketide synthase, phosphopantetheinyl transferase, *Corynebacterium glutamicum*, malonyl-CoA

## Abstract

Type I polyketide synthases (PKSs) are large multi-domain proteins converting simple acyl-CoA thioesters such as acetyl-CoA and malonyl-CoA to a large diversity of biotechnologically interesting molecules. Such multi-step reaction cascades are of particular interest for applications in engineered microbial cell factories, as the introduction of a single protein with many enzymatic activities does not require balancing of several individual enzymatic activities. However, functional introduction of type I PKSs into heterologous hosts is very challenging as the large polypeptide chains often do not fold properly. In addition, PKS usually require post-translational activation by dedicated 4’-phosphopantetheinyl transferases (PPTases). Here, we introduce an engineered *Corynebacterium glutamicum* strain as a novel microbial cell factory for type I PKS-derived products. Suitability of *C. glutamicum* for polyketide synthesis could be demonstrated by the functional introduction of the 6-methylsalicylic acid synthase ChlB1 from *Streptomyces antibioticus*. Challenges related to protein folding could be overcome by translation fusion of ChlB1_*Sa*_ to the C-terminus of the maltose-binding protein MalE from *Escherichia coli*. Surprisingly, ChlB1_*Sa*_ was also active in absence of a heterologous PPTase, which finally led to the discovery that the endogenous PPTase PptA_*Cg*_ of *C. glutamicum* can also activate ChlB1_*Sa*_. The best strain, engineered to provide increased levels of acetyl-CoA and malonyl-CoA, accumulated up to 41 mg/L (0.27 mM) 6-methylsalicylic acid within 48 h of cultivation. Further experiments showed that PptA_*Cg*_ of *C. glutamicum* can also activate nonribosomal peptide synthetases (NRPSs), rendering *C. glutamicum* a promising microbial cell factory for the production of several fine chemicals and medicinal drugs.

## Intrdouction

In microorganisms and plants, polyketide synthases (PKSs) synthesize a broad range of chemically diverse secondary metabolites, including aromatics, macrolides, polyenes, and polyethers (Shen, 2003). Due to their potent bioactivities, *e.g.* as antibiotics or antioxidants, polyketide-derived products are of interest for the development of pharmaceuticals or nutraceuticals (Kallscheuer et al., 2019; Shi et al., 2018; Yin et al., 2015). In bacteria, PKSs are typically involved in the synthesis of antibiotics or secondary lipids, whereas in plants PKSs are essential for the production of polyphenols such as stilbenes and flavonoids (Schröder, 1997).

As a common feature, PKSs catalyze the condensation of coenzyme A (CoA)-activated starter units and, in most cases, consume 3-12 malonyl-CoA molecules as building blocks during repetitive chain elongation reactions (Abe and Morita, 2010; Hertweck, 2009; Ray and Moore, 2016; Shen, 2003). This assembly process is often accompanied by defined β-keto processing reactions. The resulting intermediate is subsequently folded into the final product by an intrinsic cyclase activity of the PKS or, alternatively, subject to hydrolysis or lactonization. Once released from the PKS, the polyketide can be further modified by decorating enzymes such as methyltransferases or glycosyltransferases or it serves as a starter molecule for the synthesis of more complex compounds (Schmidlin et al., 2008; Wang et al., 2014). The architecture of PKSs is reflected by the presence of different catalytically active domains responsible for substrate selection, chain elongation, reductive processing and product release. Of particular interest are iterative type I PKSs, in which the catalytically active domains are combined in a single polypeptide chain. These domains bear acyltransferase (AT), ketosynthase (KS), ketoreductase (KR), dehydratase (DH), enoylreductase (ER) and thioesterase (TE) activities (Cox, 2007; Kage et al., 2015). During polyketide synthesis, the growing carbon chain is bound to an acyl carrier protein (ACP) domain, which must have undergone a post-translational modification to provide an active thiol group for the required thioesterification (Hertweck, 2009). This activation step is catalyzed by discrete 4’-phosphopantetheinyl transferases (PPTases), which transfer a 4’-phosphopantetheine residue from CoA to a conserved serine residue in the ACP domain, thereby converting the inactive *apo* form of the PKS to the active *holo*-PKS (Lambalot et al., 1996).

In recent years, an increasing interest in using PKSs for the production of high value compounds in heterologous microorganisms arose. However, functional expression of genes coding for type I PKSs in such hosts is challenging (Yuzawa et al., 2012). In particular, correct folding of PKS enzymes characterized by a typical length ranging from 1,500 to 4,000 amino acids and post-translational phosphopantetheinylation were identified as key issues. Nevertheless, functional introduction of several PKSs was achieved *e.g.* in *Escherichia coli* and *Streptomyces* spp. by using strategies for improved protein folding and by co-expression of PPTase-encoding genes (Baltz, 2010; Liu et al., 2015; Ugai et al., 2015).

Since decades, *Corynebacterium glutamicum* is used at industrial scale for the production of amino acids, in particular of L-glutamate and L-lysine (Eggeling and Bott, 2015), but this bacterium was also engineered for the synthesis of other biotechnologically interesting compounds, *e.g.* alcohols, diamines, dicarboxylic acids, aromatic compounds and secondary metabolites (Heider et al., 2014; Kallscheuer and Marienhagen, 2018; Kallscheuer et al., 2016a; Litsanov et al., 2012; Nguyen et al., 2015; Vogt et al., 2016). In previous studies, we already established the functional introduction of plant-derived type III PKSs with a typically length of 300-400 amino acids into *C. glutamicum*, which enabled plant polyphenol synthesis with this bacterium (Kallscheuer et al., 2016b; Milke et al., 2019b).

In this study, we introduce *C. glutamicum* as a promising microbial host for polyketide production as we could functionally introduce the type I PKS 6-methylsalicylic acid synthase ChlB1 from *Streptomyces antibioticus* (ChlB1_*Sa*_, 1,756 aa, 186 kDa, UniProt ID Q0R4P8) into this bacterium (Jia et al., 2006; Shao et al., 2006). ChlB1_*Sa*_ catalyzes the conversion of the starter unit acetyl-CoA and three molecules of malonyl-CoA into the aromatic compound 6-methylsalicylic acid (6-MSA) (Fig. 1) (Parascandolo et al., 2016). The latter is an important building block in the biosynthesis of several antibiotics, including chlorothricin, maduropeptin, pactamycin, and polyketomycin (Daum et al., 2009; Ito et al., 2009; Jia et al., 2006; Van Lanen et al., 2007).

**Figure 1.**
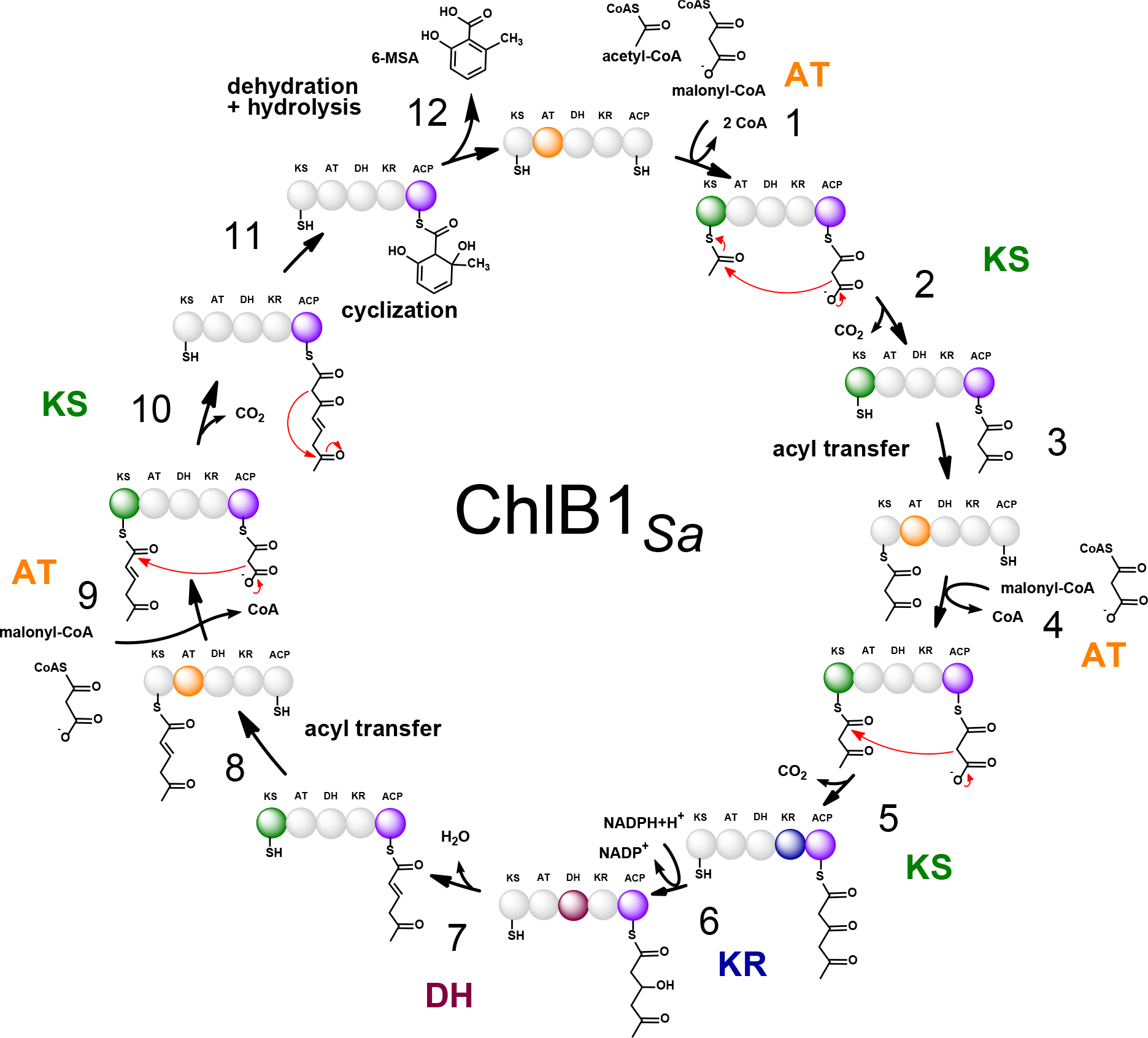
6-Methylsalicylic acid formation by the type I PKS ChlB1. The depicted steps follow a previously established reaction sequence. 6-Methylsalicylic acid is produced from acetyl-CoA and three molecules of malonyl-CoA. Acetyl-CoA as starter unit is consumed in step 1, whereas three malonyl-CoA molecules are consumed in steps 1, 4, and 9, respectively. The required domains involved in the respective reaction steps are highlighted by different colors. ACP: acyl carrier protein domain, AT: acyltransferase domain, KS: ketosynthase domain, KR: ketoreductase domain, DH: dehydratase domain, 6-MSA: 6-methylsalicylic acid.

## Material and methods

### Bacterial strains, plasmids, media and growth conditions

All bacterial strains and plasmids used in this study and their relevant characteristics are listed in Table 1. *C. glutamicum* was routinely cultivated aerobically at 30 °C in brain heart infusion (BHI) medium (Difco Laboratories, Detroit, USA) or in defined CGXII medium with 4% (w/v) glucose as sole carbon and energy source (Keilhauer et al., 1993). *E. coli* DH5α used for plasmid constructions was cultivated in LB medium (Bertani, 1951) at 37 °C. For maintenance of plasmids, kanamycin (50 μg/mL for *E. coli* or 25 μg/mL for *C. glutamicum*), chloramphenicol (25 μg/mL for *E. coli*) or spectinomycin (100 μg/mL for *E. coli* and *C. glutamicum*) was added to the medium. Bacterial growth was followed by measuring the optical density at 600 nm (OD_600_). *C. glutamicum* was grown for 6-8 hours in test tubes with 5 mL BHI medium on a rotary shaker at 170 rpm (first preculture) and was subsequently inoculated into 50 mL defined CGXII medium with 4 % (w/v) glucose in 500 mL baffled Erlenmeyer flasks (second preculture). Cell suspensions were cultivated overnight on a rotary shaker at 130 rpm. The main culture was inoculated to an OD_600_ of 5 in defined CGXII medium with 4 % (w/v) glucose and heterologous gene expression was induced one hour after inoculation using the indicated amount of IPTG.

**Table 1.**
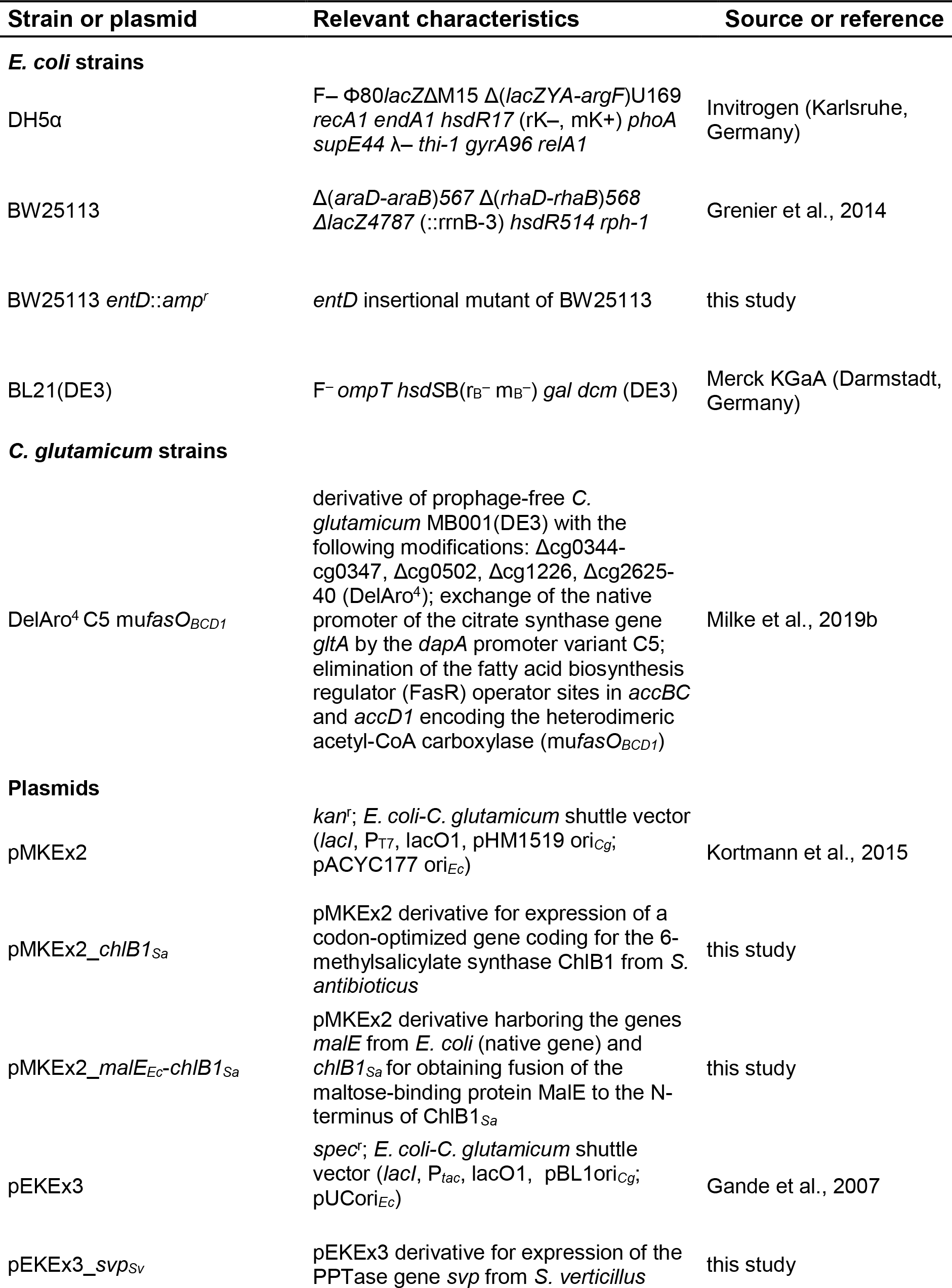
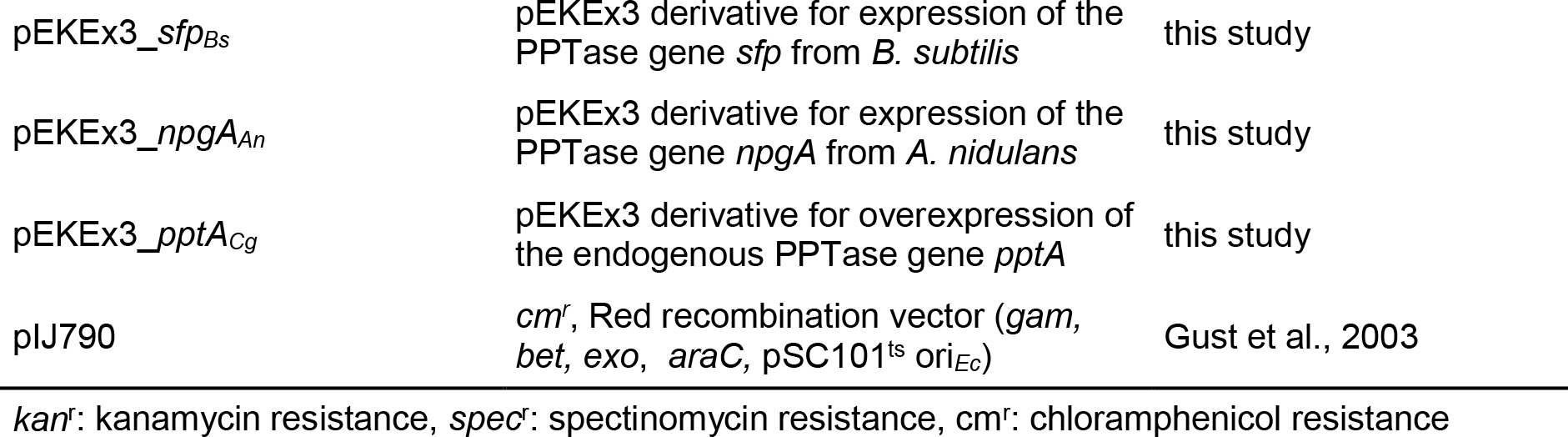
Strains and plasmids used in this study.

### Construction of plasmids and strains

Plasmids and strains used in this study are listed in table 1. Standard protocols of molecular cloning, such as PCR, DNA restriction, and ligation (Sambrook and Russell, 2001), were carried out for recombinant DNA work. Techniques specific for *C. glutamicum*, *e.g.* electroporation for transformation of strains, were performed as described previously (Eggeling and Bott, 2005). Synthetic genes were obtained from Thermo Fisher Scientific, formerly GeneArt (Darmstadt, Germany). Genes were amplified by PCR using oligonucleotides with unique restriction sites for cloning (Table 2). All constructed plasmids were finally verified by DNA sequencing at Eurofins Genomics (Ebersberg, Germany). The PPTase gene *entD* in *E. coli* BW25113 was inactivated by λ/Red recombineering (Datsenko and Wanner, 2000). For this purpose, the ampicillin resistance gene from pUC19 was furnished with *entD* homologous arms by PCR using oligonucleotides *entD*-s and *entD*-as. The 1.3 kb-sized amplicon was introduced into *E. coli* BW25113/pIJ790 by electroporation after induction of λ *red* genes following a previously reported procedure (Kreutzer et al., 2011). The resulting transformants were incubated overnight at 37 °C on LB agar containing ampicillin (50 μg/mL). To identify recombinants with the desired mutation, PCRs using oligonucelotides *entD*SQ-s and *entD*SQ-as were conducted. The mutation in *E. coli* BW25113 *entD*::*amp*^r^ was confirmed by DNA sequencing.

**Table 2.**
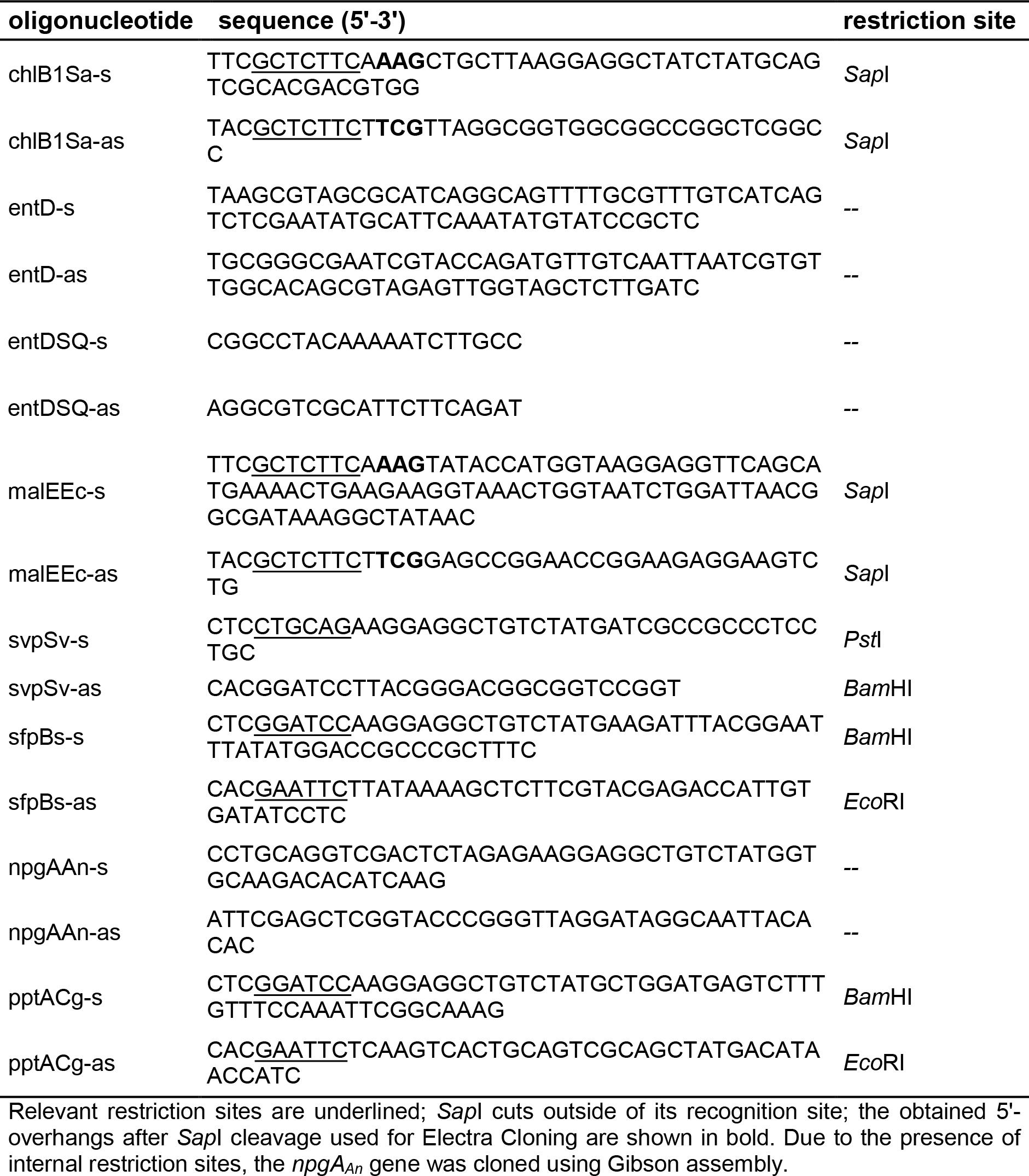
Oligonucleotides used in this study.

### LC-MS analysis for quantification of 6-MSA in culture supernatants

6-MSA was quantified in culture supernatants by LC-MS using an Agilent ultra-high-erformance LC 1290 Infinity System coupled to a 6130 Quadrupole LC-MS System (Waldbronn, Germany). LC separation was carried out using a Kinetex 1.7u C_18_ 100 Å pore size column (50 mm by 2.1 mm (internal diameter)) (Phenomenex, Torrance, CA, USA) at 50°C. For elution, 0.1 % (v/v) acetic acid (solvent A) and acetonitrile supplemented with 0.1 % (v/v) acetic acid (solvent B) were applied as the mobile phases at a flow rate of 0.3 mL/min. A gradient was used, where the amount of solvent B was increased stepwise: minute 0 to 6: 5% to 30 %, minute 6 to 7: 30 % to 50 %, minute 7 to 8: 50 % to 100 % and minute 8 to 8.5: 100 % to 5 %. The mass spectrometer was operated in the negative electrospray ionization (ESI) mode, and data acquisition was performed in selected-ion-monitoring (SIM) mode. Authentic metabolite standards were purchased from Thermo Fisher Scientific, formerly Acros Chemicals (Geel, Belgium). Area values for [M-H]^-^ mass signals were linear up to metabolite concentrations of at least 250 mg/L. Benzoic acid (final concentration 100 mg/L) was used as internal standard. Calibration curves were calculated based on analyte/internal standard ratios for the obtained area values.

### PPTase complementation in an *entD*-deficient *E. coli* variant

The promiscuity of PptA_*Cg*_ was tested through the restoration of enterobactin biosynthesis by complementing the inactivated *entD* gene. For this, *E. coli* BW25113 *entD*::*amp*^r^ was transformed either with the empty vector pEKEx3 or with the vector pEKEx3_*pptA*_*Cg*_ harboring the *pptA*_*Cg*_ gene from *C. glutamicum*. For analysis of enterobactin production, the two resulting *E. coli* strains were cultivated in 50 mL M9 mineral medium without iron supplementation at 37 °C to an OD_600_ of 0.6. At this time, expression of the *pptA*_*Cg*_ gene was induced with 1 mM IPTG. Afterwards, the incubation was continued overnight at 37°C. To verify the production of enterobactin, the cultures were acidified with hydrochloric acid (pH 2) and exhaustively extracted with ethyl acetate. After removal of the organic solvent, the crude extracts were dissolved in methanol. The extracts were analyzed by HPLC-MS using an Agilent 1260 Infinity System (Agilent, Waldbronn, Germany) coupled to a Bruker Compact ESI-Q-TOF mass spectrometer (Bruker, Bremen, Germany) equipped with a Nucleoshell RP-C18 column (100 mm × 2 mm, 2.7 μm, Macherey Nagel, Düren, Germany). A linear gradient of acetonitrile in water with 0.1 % (v/v) formic acid (from 5% to 98% acetonitrile within 10 min; flow rate, 0.4 mL/min) was used for metabolic profiling. UV chromatograms were recorded at 316 nm.

## Results

### Functional introduction of *chlB1_Sa_* into *C. glutamicum* enables 6-MSA synthesis

In the course of developing a *C. glutamicum* platform strain for plant polyphenol synthesis, the central carbon metabolism was reengineered towards increased availability of the PKS substrates acetyl-CoA and malonyl-CoA, and the entire catabolic network for aromatic compounds was eliminated to avoid any product degradation (Kallscheuer et al., 2016b; Milke et al., 2019a; Milke et al., 2019b). Against this background, *C. glutamicum* DelAro^4^ C5 mu*fasO*_*BCD1*_ characterized by (I) deletion of 21 genes involved in the degradation of various aromatic compounds, (II) reduction of the citrate synthase activity and (III) deregulation of genes encoding the acetyl-CoA carboxylase, was selected as parent strain for further experiments (Fig. 2). Initially, it was tested if *C. glutamicum* DelAro^4^ C5 mu*fasO*_*BCD1*_ can metabolize supplemented 6-MSA. However, cultivations in the presence of 200 mg/L (1.3 mM) 6-MSA and analysis of culture supernatants by LC-MS confirmed that *C. glutamicum* DelAro^4^ C5 mu*fasO*_*BCD1*_ is neither capable of catabolizing nor unspecifically modifying this compound (data not shown).

**Figure 2.**
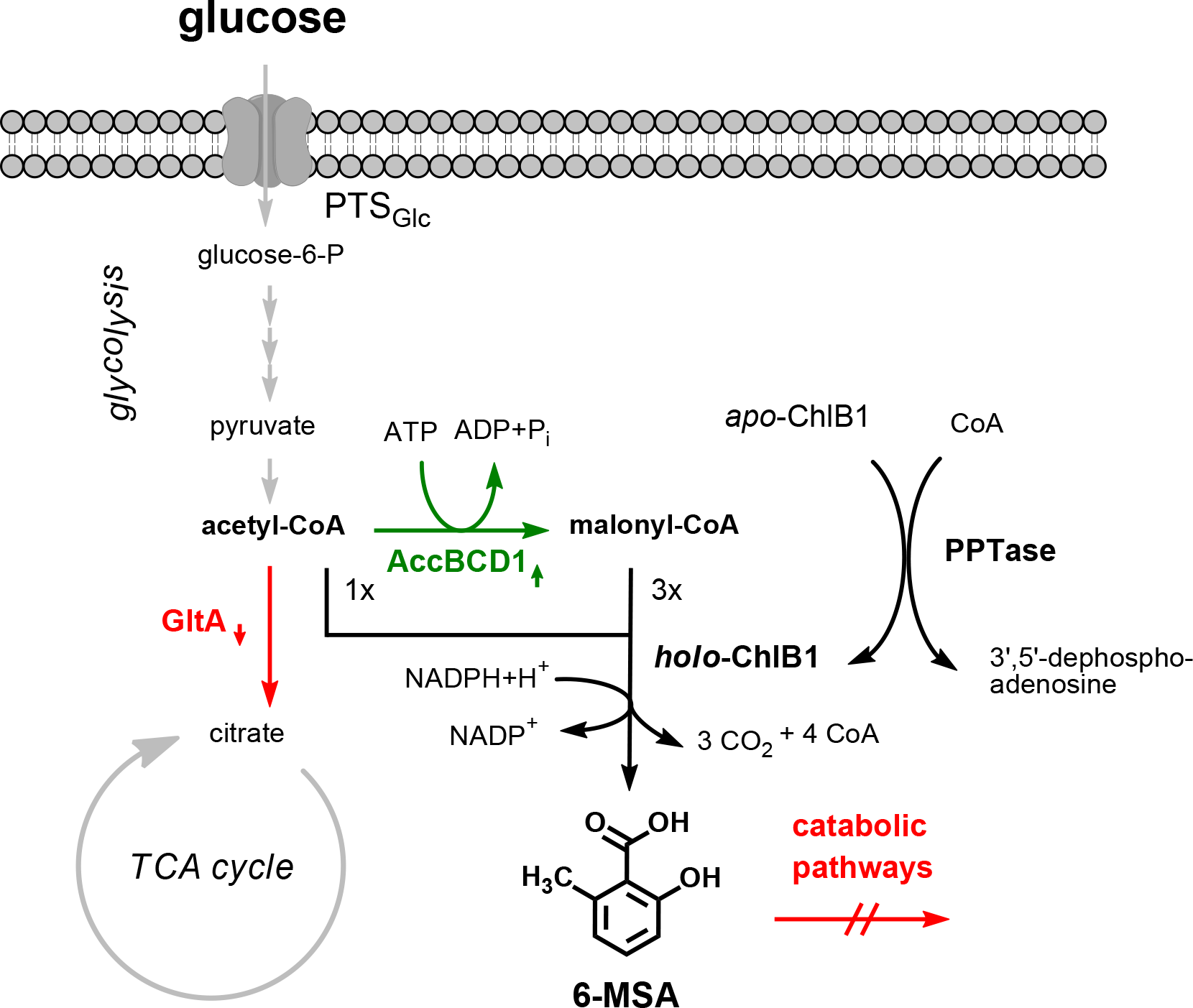
Metabolic engineering of *C. glutamicum* for 6-MSA production. The engineered central carbon metabolism of *C. glutamicum* and the ChlB1 and PPTase-catalyzed reactions are shown. During the construction of *C. glutamicum* DelAro^4^ C5 mu*fasO*_*BCD1*_ used for 6-MSA production, the following genomic modifications were introduced: (I) Replacement of the native *gltA* promoter by the constitutive *dapA* promoter variant C5 for reducing the overall citrate synthase activity (GltA) activity to 5 % (compared to the GltA activity in the *C. glutamicum* wild type); (II) mutation of regulator binding sites for the transcriptional repressor FasR upstream of the open reading frames of *accBC* and *accD1* (coding for the heterodimeric acetyl-CoA carboxylase AccBCD1) to abolish gene repression; (III) deletion of genes coding for enzymes involved in the degradation of aromatic compounds to avoid any potential 6-MSA consumption by *C. glutamicum*. PTSGlc: glucose-specific phosphotransferase system, PPTase: 4’-phosphopantetheinyl transferase, TCA cycle: tricarboxylic acid cycle.

For heterologous expression of the 6-MSA synthase gene from *S. antibioticus*, the codon-optimized *chlB1*_*Sa*_ gene was cloned into the expression plasmid pMKEx2, which allows for an IPTG-inducible gene expression under the control of the strong T7 promoter (Kortmann et al., 2015). In addition, a gene coding for the broad-spectrum PPTase Svp from *Streptomyces verticillus* (Svp_*Sv*_, UniProt ID Q9F0Q6) was co-expressed from the plasmid pEKEx3_*svp*_*Sv*_ for phosphopantetheinylation of ChlB1_*Sa*_ (Sánchez et al., 2001). Unfortunately, the constructed strain *C. glutamicum* DelAro^4^ C5 mu*fasO*_*BCD1*_ pMKEx2_*chlB1*_*Sa*_ pEKEx3_*svp*_*Sv*_ failed to produce any LC-MS-detectable amounts of 6-MSA when cultivated in defined CGXII medium with 4 % (w/v) glucose and two different IPTG concentrations (20 μM or 1 mM). However, problems related to folding of “challenging” proteins such as ChlB1_*Sa*_ were not unexpected when considering its protein size of 186 kDa. Thus, we decided to test translational fusion of ChlB1_*Sa*_ to the C-terminus of the maltose binding protein MalE of *E. coli*. Although being even larger than ChlB1_*Sa*_ (1,756 aa, 186 kDa), the MalE_*Ec*_-ChlB1_*Sa*_ fusion protein (2,139 aa, 228 kDa) appeared to be active in *C. glutamicum* as accumulation of 2 mg/L (0.013 mM) and 6 mg/L 6-MSA (0.039 mM) after induction with 20 μM or 1 mM IPTG, respectively, could be detected (Fig. 3a,b). Neither under non-inducing conditions (no IPTG) nor in case of cultivation of a control strain harboring only the two empty plasmids, 6-MSA synthesis could be observed.

**Figure 3.**
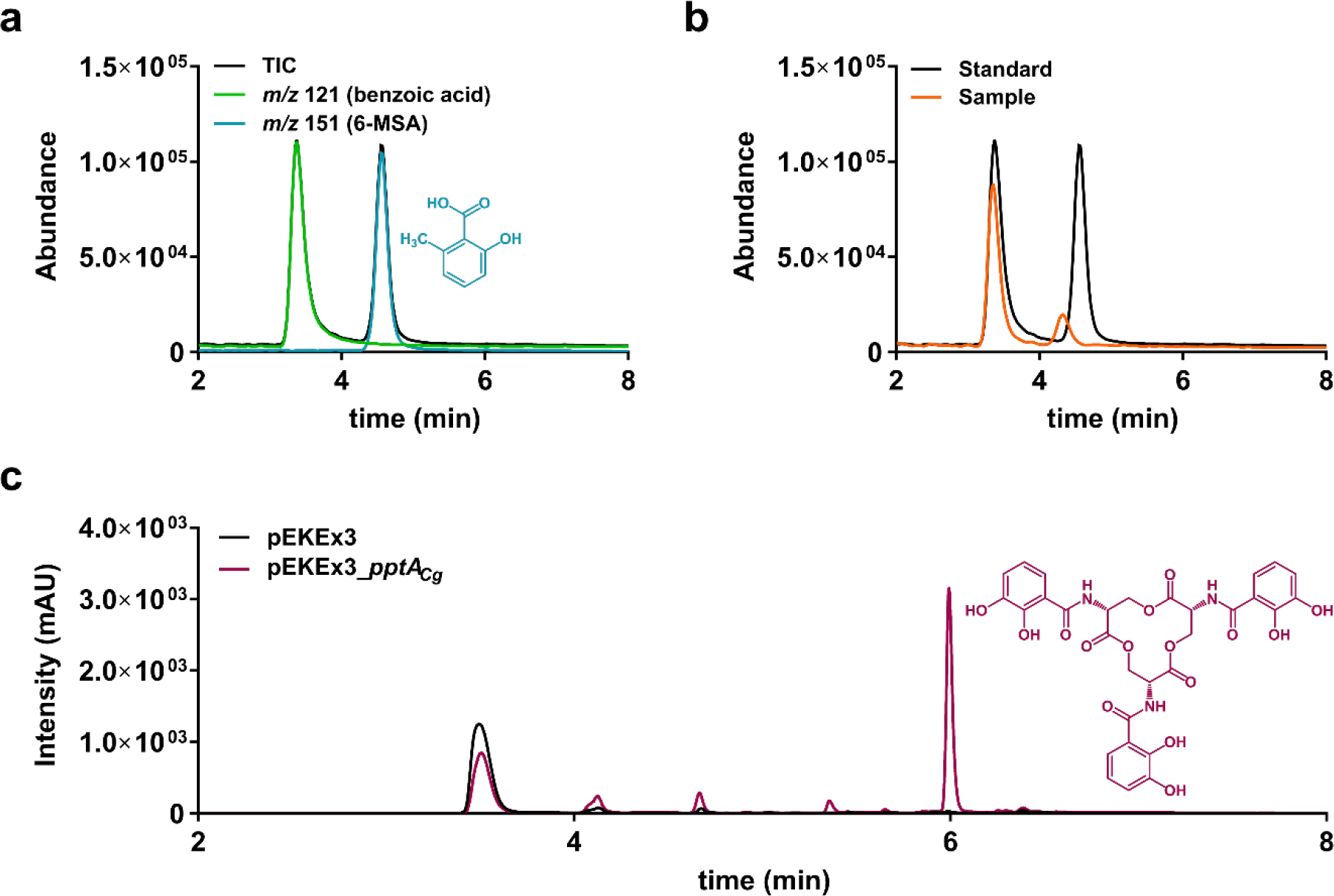
Analysis of 6-MSA production with *C. glutamicum* and enterobactin synthesis in *E. coli*. **(a)**Chromatogram of an authentic 6-MSA/benzoic acid standard, both as total ion current and as individual *m/z* ratios for 6-MSA and benzoic acid, respectively. **(b)**Total ion current chromatograms of a typical sample obtained from the cultivation of *C. glutamicum* DelAro^4^ C5 mu*fasOBCD1* pMKEx2_*malEEc*-*chlB1Sa* pEKEx3_*svpSv* producing 6-MSA and an authentic 6-MSA standard for comparison. A slight shift in the retention time of the 6-MSA peak in culture supernatants of *C. glutamicum* in comparison to the metabolite standard was observed. Integrity of the shifted 6-MSA peak was verified by addition of 6-MSA to the culture supernatant (spiking). **(c)**UV-chromatograms of culture supernatants from *E. coli* BW25113 *entD*::*amp*^r^ harboring either an empty plasmid (pEKEx3) or a plasmid featuring *pptA_Cg_* from *C. glutamicum* (pEKEx3_*pptA_Cg_*). Both chromatograms were recorded at 316 nm. The enterobactin peak at a retention time of 6 min was identified by ESI-MS analysis.

Formation of 6-MSA suggested that ChlB1_*Sa*_ can be successfully activated by the PPTase Svp_*Sv*_. However, the *svp_Sv_*-expressing *C. glutamicum* strain only reached a drastically reduced final biomass (OD_600_= 30 - 33) during the cultivation and 6-MSA production experiments. Considering that standard CGXII medium contains only 1 mM MgSO_4_ as sole source of Mg^2+^-ions required as metal cofactor for PPTase activity, we assumed that heterologous expression of *svp*_*Sv*_ leads to rapid depletion of Mg^2+^ ions, which in turn would limit overall cell growth. Hence, different MgSO_4_ concentrations ranging from 15 – 200 mM were tested in cultivations of the engineered *C. glutamicum* strain (Fig. 4). These experiments revealed, that limited Mg^2+^ availability was indeed the reason for the observed reduced-growth-phenotype since all cultures with MgSO_4_ concentrations > 1 mM reached a final OD_600_ between 50 to 55, typical for shake flask cultivations of *C. glutamicum*. In this context, it was not surprising that low Mg^2+^ concentrations also limited 6-MSA synthesis as cultivations with 15 mM MgSO_4_ doubled product titers, reaching 14 mg/L (0.09 mM) 6-MSA. The highest product titer of 20 mg/L (0.13 mM) at this stage was determined in cultivations with supplementation of 50 mM MgSO_4_, whereas supplementation of 100 mM MgSO_4_ and above had a negative effect on product formation (Fig. 3). All subsequent cultivations for 6-MSA production were thus performed in defined CGXII medium containing 50 mM MgSO_4_.

**Figure 4.**
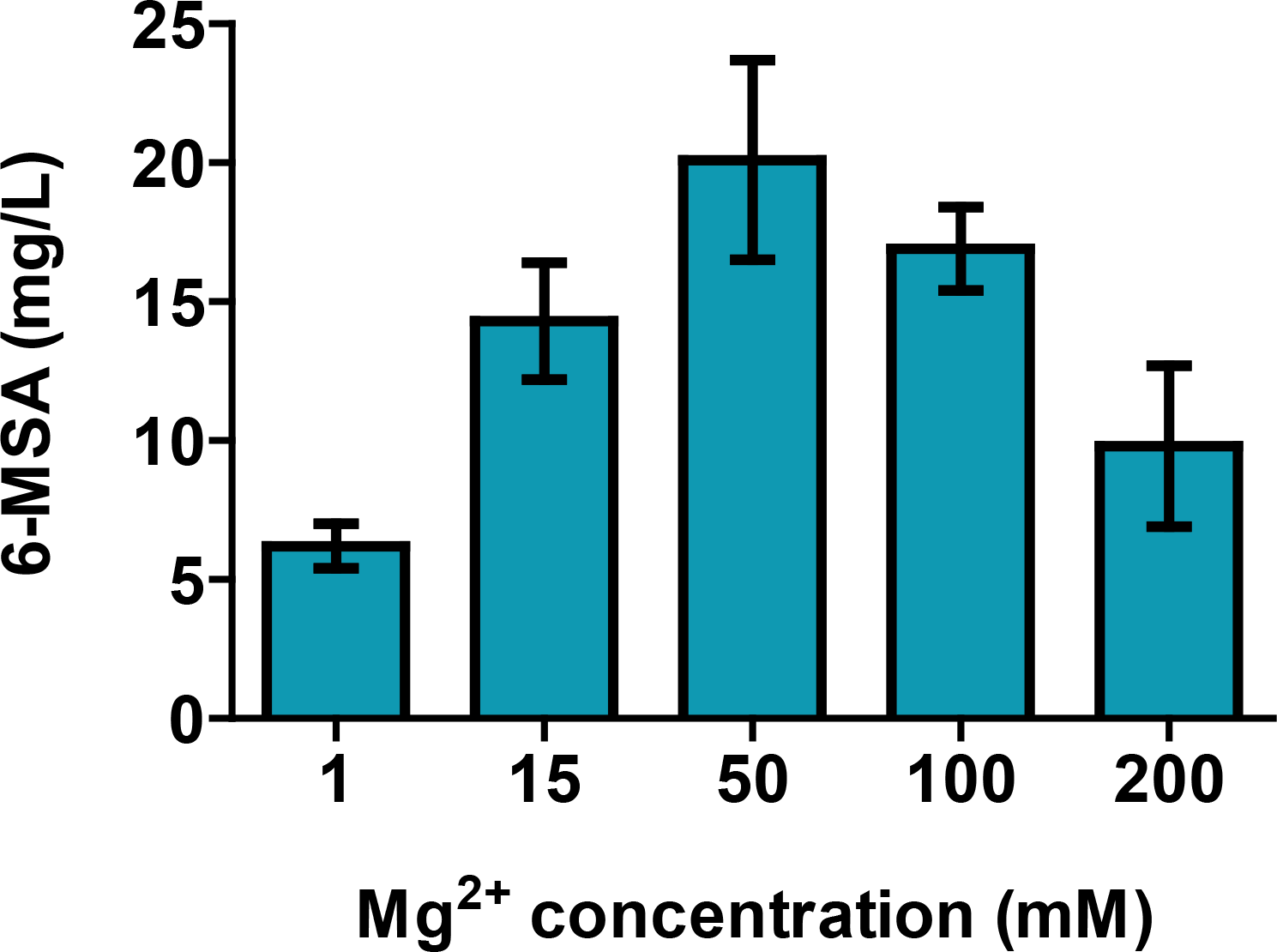
Variation of the Mg^2+^ concentration in the culture medium for optimizing PPTase activity in *C. glutamicum*. MgSO_4_ concentrations ranging from 1 to 200 mM were tested during the microbial production of 6-MSA in defined CGXII medium with 4 % glucose, and 1 mM IPTG using *C. glutamicum* DelAro^4^ C5 mu*fasOBCD1* pMKEx2_*chlB1Sa* pEKEx3_*svpSv*. Notably, the standard MgSO_4_ concentration in defined CGXII medium is 1 mM (first column). Data represent average values and standard deviations of three biological replicates.

### An endogenous PPTase of *C. glutamicum* can activate ChlB1_*Sa*_

In microbial host systems engineered for polyketide synthesis involving type I PKS, PPTase-mediated PKS activation is essential for product formation, but not every PPTase is capable of activating every PKS (Cox et al., 1997; Shen et al., 1992). The PPTase Svp_*Sv*_ from *S. verticillus* was initially selected due to its well-described ability to activate a broad range of different type I PKSs and the close relationship of *S. verticillus* to *S. antibioticus*, from which ChlB1_*Sa*_ originates (Sánchez et al., 2001). With the aim to identify an even more suitable PPTase for ChlB1_*Sa*_ activation in *C. glutamicum*, we evaluated the performance of other broad-spectrum PPTases such as Sfp_*Bs*_ (UniProt ID P39135) from *Bacillus subtilis* and NpgA_*An*_ (UniProt ID G5EB87) from *Aspergillus nidulans* (Oberegger et al., 2003; Quadri et al., 1998). Interestingly, *C. glutamicum* DelAro^4^ C5 mu*fasO*_*BCD1*_ pMKEx2_*malE_Ec_*-*chlB1*Sa pEKEx3_*npgA*_*An*_ expressing the gene for the PPTase from *A. nidulans* did not grow at all and thus also failed to produce 6-MSA. Since this was not further investigated we can only speculate that the *npgA*_*An*_ gene product is toxic for *C. glutamicum.* In contrast, coexpression of *sfp*_*Bs*_ enabled accumulation of 28 mg/L (0.18 mM) 6-MSA. To our surprise, the highest product titer was obtained in cultivations of a constructed *C. glutamicum* DelAro^4^ C5 mu*fasO*_*BCD1*_ variant without any heterologous expression of a PPTase-encoding gene, which was originally supposed to serve as negative control. This strain, *C. glutamicum* DelAro^4^ C5 mu*fasO*_*BCD1*_ pMKEx2_*malE_Ec_*-*chlB1_Sa_*, accumulated 41 mg/L (0.27 mM) 6-MSA, indicating that *C. glutamicum* harbors an endogenous enzyme capable of activating ChlB1_*Sa*_. At this point, we speculated that this endogenous PPTase activity is present in all constructed strains and that lower 6-MSA concentrations determined in cultivations of *C. glutamicum* strains expressing heterologous PPTase genes are due to the metabolic burden of the plasmid-based expression of additional genes from a second plasmid also requiring supplementation of an additional antibiotic. Analysis of the genome sequence of *C. glutamicum* ATCC 13032 identified two genes encoding PPTases, namely *pptA* (cg2171) and *acpS* (cg2738). AcpS_*Cg*_ (UniProt ID Q8NMS4) was shown to be crucial for the activation of the fatty acid synthases FasI-A and FasI-B in this bacterium (Chalut et al., 2006). The putative function of PptA_*Cg*_ (UniProt ID Q8NP45) is the activation of the sole type I PKS13 in *C. glutamicum* (Cg3178, 1610 aa, 172 kDa), which is required for the synthesis of corynomycolic acids as important building blocks for cell wall synthesis in coryneform bacteria (Gande et al., 2004).

Motivated by the finding that an endogenous PPTase of *C. glutamicum* is capable of activating ChlB1_*Sa*_, we compared the sequence of the two endogenous PPTase candidates PptA_*Cg*_ and AcpS_*Cg*_ to Svp_*Sv*_. PptA_*Cg*_ and Svp_*Sv*_ share a sequence identity of 34 % (sequence similarity: 58 %), whereas AcpS_*Cg*_ appears to be unrelated to Svp_*Sv*_ (sequence identity: 8 %; sequence similarity: 19 %) (Fig. S1). Hence, it appears to be more likely that PptA_*Cg*_ rather than AcpS_*Cg*_ can activate ChlB1_*Sa*_ in *C. glutamicum*. This notion is also supported by the domain architecture as both enzymes, PptA_*Cg*_ and Svp_*Sv*_, belong to the EntD superfamily of PPTases (NCBI domain cl27525), whereas AcpS_*Cg*_ is a PPTase of the ACPS superfamily (NCBI domain cl00500). In order to confirm the relevance of PptA_*Cg*_ for ChlB1_*Sa*_ activation, deletion of the respective gene *pptA* was planned, However, a *pptA* deletion mutant was previously shown to exhibit a severe growth phenotype reflected in cell aggregation during cultivation in liquid medium (Chalut et al., 2006). For this reason, the idea of deleting *pptA* was abandoned as discrimination between lacking 6-MSA production due to missing PptA_*Cg*_-mediated ChlB1_*Sa*_-activation or poor growth is not possible. Instead, it was tested whether overexpression of *pptA* could further improve 6-MSA production with *C. glutamicum*. The strain *C. glutamicum* DelAro^4^ C5 mu*fasO*_*BCD1*_ pMKEx2_*malE_Ec_*-*chlB1*_*Sa*_ pEKEx3_*pptA*_*Cg*_ accumulated only 18 mg/L (0.12 mM) 6-MSA, which further supports the notion that reduced 6-MSA titers in the strains additionally expressing PPTase-encoding genes are due to an increased metabolic burden and that the native *pptA*_*Cg*_ expression is already sufficient for ChlB1_*Sa*_ activation.

### PptA of *C. glutamicum* is a broad-spectrum PPTase

To interrogate a possible broad-spectrum PPTase activity of PptA_*Cg*_, further experiments were performed in *E. coli*. This bacterium is in general capable of producing the catecholate siderophore enterobactin under iron-limited conditions (Grass, 2006). Biosynthesis of enterobactin involves six proteins of which two, EntB and EntF, feature carrier protein domains (Gehring et al., 1998). Analogous to PKS systems, both EntB and EntF are phosphopantetheinylated by a PPTase named EntD (UniProt ID P19925) encoded in the enterobactin locus (Gehring et al., 1997; Lambalot et al., 1996). While inactivation of *entD* abolishes production of enterobactin in *E. coli* (Cox et al., 1970), this phenotype can be rescued by complementation with an exogenous PPTase (Barekzi et al., 2004). To test whether PptA_*Cg*_ is similarly capable to act as a substitute for the enterobactin PPTase, an *entD* mutation was introduced into the *E. coli* strain BW25113, which was previously reported as enterobactin producer (Ma and Payne, 2012). Comparative LC-MS analyses with the wild type confirmed the loss of enterobactin in the *E. coli* BW25113 *entD*::*amp*^r^ strain (data not shown). Hence, it was evident that *E. coli* BW25113 does not possess another PPTase, which could compensate for the functional loss of EntD. Following the expression of *pptA*_*Cg*_ in *E. coli* BW25113 *entD*::*amp*^r^, the mutant resumed the production of enterobactin, which strongly suggests that PptA_*Cg*_ is capable to phosphopantetheinylate the two carrier protein domains in EntB and EntF (Fig. 3c). This experiment indicates that PptA_*Cg*_ of *C. glutamicum* can indeed activate various carrier protein domains and can thus be regarded as broad-spectrum PPTase with many possible applications in microbial natural product synthesis.

## Discussion

Iterative type I PKSs biocatalysts are particularly appealing for the production of high-value molecules from simple CoA-activated precursors derived from the central carbon metabolism. Once an iterative PKS is activated, all reaction steps are catalyzed by individual domains, in that sense the PKS itself represents a biochemical “assembly line” within a microbial cell factory. This avoids to a large degree any engineered deregulation of natural biosynthetic pathways and thereby circumvents the accumulation of undesired side-products. In consequence, iterative PKSs can be considered as ideal targets for the production of chemical building blocks as well as medicinal drugs in engineered microbial cell factories.

Here, we functionally integrated the 6-MSA synthase ChlB1 from *S. antibioticus* into *C. glutamicum* and demonstrated its suitability as a cell factory for 6-MSA synthesis. Production of related hydroxybenzoic acids in engineered microorganism such as salicylic acid (a derivative of 6-MSA lacking the methyl group) requires deregulation and engineering of the shikimate pathway and can lead to metabolic imbalances, auxotrophic strains or undesired accumulation of side products (Kallscheuer and Marienhagen, 2018; Lin et al., 2014). In contrast, after introduction of ChlB1_*Sa*_ in *C. glutamicum* no significant accumulation of side products could be observed.

Microbial production of 6-MSA using the 6-MSA synthase from *Penicillium patulum* was already demonstrated earlier using *E. coli* and *Saccharomyces cerevisiae* as production hosts (Kealey et al., 1998; Wattanachaisaereekul et al., 2007; Wattanachaisaereekul et al., 2008). For the essential PPTase-mediated activation of the heterologous PKS, co-expression of a broad spectrum PPTase gene, such as *sfp* from *B. subtilis*, was mandatory in both hosts. In the direct comparison, *S. cerevisiae* was more suitable for 6-MSA production as titers from 0.2-1.7 g/L (1.3-11.2 mM) 6-MSA were reported, whereas *E. coli* accumulated only 75 mg/L (0.49 mM) (Kealey et al., 1998; Wattanachaisaereekul et al., 2008). The determined product concentrations in *E. coli* are comparable to 6-MSA titers obtained using engineered *Streptomyces coelicolor* strains (Bedford et al., 1995) and with *C. glutamicum* in this study. The performance of the different production hosts, however, is difficult to compare as we decided to use ChlB1 from *S. antibioticus* in this study instead of the enzyme from *P. patulum*.

In both organisms, *C. glutamicum* and *E. coli*, challenges related to protein insolubility needed to be addressed. In our study improved folding was achieved by translation fusion of the PKS to the maltose-binding protein MalE, whereas co-expression of chaperone-encoding genes from *S. coelicolor* improved folding and solubility of different type I PKSs in *E. coli* (Betancor et al., 2008). In case of the essential PPTase-mediated activation of PKSs, we could show that *C. glutamicum* harbors a broad-spectrum PPTase (PptA_*Cg*_), which is not only capable of activating ChlB1_*Sa*_, but also activates the enterobactin biosynthesis enzymes EntB and EntF in *E. coli*. It is thus likely that expression of a heterologous PPTase gene can be omitted in future PKS- and NRPS-related applications using *C. glutamicum*, which will significantly simplify and shorten strain development times as only the core biosynthesis gene needs to be functionally expressed. This important property of *C. glutamicum* cannot be valued highly enough as phosphopantetheinylation in heterologous hosts often fails due to the inavailability of a suitable PPTase (Crosby et al., 1995; Li and Neubauer, 2014; Roberts et al., 1993; Shen et al., 1992).

Taken together, we consider *C. glutamicum* as a promising host organism for microbial polyketide production using type I PKSs. *C. glutamicum* is an actinomycete and thus closely related to Streptomycetes, which are a valuable source of many interesting type I PKSs (including ChlB1_*Sa*_). Access to *C. glutamicum* variants with increased acetyl-CoA and malonyl-CoA availability and easy-to-handle production plasmids enabling the expression of soluble MalE-fusion proteins further underline the potential of this microorganism for polyketide synthesis.

