## Supplemental Information for "Competence of *Corynebacterium glutamicum* as a host for the production of type I polyketides"

*ORCID-IDs:*
Nicolai Kallscheuer (0000-0003-4925-6923)
Lars Milke (0000-0001-9151-1065)
Markus Nett (0000-0003-0847-086X)
Jan Marienhagen (0000-0001-5513-3730)

**Running Title:** 6-methylsalicylate production in *C. glutamicum*

**Keywords:** polyketide, 6-methylsalicylate, type I polyketide synthase, phosphopantetheinyl transferase, *Corynebacterium glutamicum*, malonyl-CoA

**a**

```
PptA(Cg)      1    MLDESLFPNSAK--FSFIKTGDAVNLDHFHQLHPLEKALVAHSVDIRKAEFGDARWCAHQ   58
Svp(Sv)       1    -MIAALLPSWAVTEHAFTD----APDDPVSLLFPEEAAHVARAVPKRLHEFATVRVCARA   55
                    :  :*:*. *   .:* .    .  * .  *.* * * **::*  *  **. .* **:

PptA(Cg)     59    ALQALGRDSGDPILRGERGMPLWPSSVSGSLTHTDGFRAAVVAPRLLVRSMGLDAEPAEP  118
Svp(Sv)      56    ALGRLGLPP-GPLLPGRRGAPSWPDGVVGSMTHCQGFRGAAVARAADAASLGIDAEPNGP  114
                   **  **    .*:* *.** * **..* **:** :***.*.**    . *:*:****  *

PptA(Cg)    119    LPKDVLGSIARVGEIPQLKRLEEQ-GVHCADRLLFCAKEATYKAWFPLTHRWLGFEQAEI  177
Svp(Sv)     115    LPDGVLAMVSLPSEREWLAGLAARRPDVHWDRLLFSAKESVFKAWYPLTGLELDFDEAEL  174
                   **..**. ::  .*   *  *  :      *****.***:.:***:***   *.*::**:

PptA(Cg)    178    DLRDDG-TFVSYLLVRPTP------VPFISGKWVLRDGYVIAATAVT-------------  217
Svp(Sv)     175    AVDPDAGTFTARLL-VPGPVVGGRRLDGFEGRWAAGEGLVVTAIAVAAPAGTAEESAEGA  233
                    :  *. **.: **  * *      :  :.*:*.  :* *::* **:

PptA(Cg)    218    -------------                                                 217
Svp(Sv)     234    GKEATADDRTAVP                                                 246
```

**b**

```
AcpS(Cg)      1    ------------------------------------------------------------    0
Svp(Sv)       1    MIAALLPSWAVTEHAFTDAPDDPVSLLFPEEAAHVARAVPKRLHEFATVRVCARAALGRL   60

AcpS(Cg)      1    ------------------------------------------------------------    0
Svp(Sv)      61    GLPPGPLLPGRRGAPSWPDGVVGSMTHCQGFRGAAVARAADAASLGIDAEPNGPLPDGVL  120

AcpS(Cg)      1    -MISIGTDLVHI------------------------------------------------   11
Svp(Sv)     121    AMVSLPSEREWLAGLAARRPDVHWDRLLFSAKESVFKAWYPLTGLELDFDEAELAVDPDA  180
                    *:*: ::   :

AcpS(Cg)     12    SAFAEQLAQPGSSFMEVFSAGERRKANERQASRYAEHLAGRWAAKESFIKAWSQAIYGQP   71
Svp(Sv)     181    GTFTARLLVPGP----------------VVGGRRLDGFEGRWAAGEGLVVTAIAV--AAP  222
                   .:*: :*  **                   ..*  : : ***** *.:: :   .  . *

AcpS(Cg)     72    PVIAEEAVVWRDIEVRADAWGRVAIELAPELAAVVRESIGEFSSSLSISHDGDYAVATCV  131
Svp(Sv)     223    AGTAEESAEGAGKEATADD--RT-------------------------------AVP---  246
                      ***:.   . *. **   *.                               **

AcpS(Cg)    132    LTIQ                                                          135
Svp(Sv)     247    ----                                                          246
```

**Figure S1. Sequence alignment of different PPTases evaluated for an application in *C. glutamicum*. (a)** Sequence comparison of the native PPTase PptA of *C. glutamicum* and Svp from *Streptomyces verticillus*; **(b)** Sequence comparison of the native PPTase AcpS of *C. glutamicum* and Svp from *Streptomyces verticillus*. Alignments were performed at the UniProt database website using Clusal Omega (Sievers *et al.* 2011).
